# Dedifferentiation-Driven Oncogenic Stemness Promotes Tumor-Sustaining Adaptability in the Intestinal Epithelium

**DOI:** 10.1101/2025.10.22.683922

**Authors:** Kylee Zgeib, Thomson Hui, Simon Garcia, Zahra Hashemi, Shima Nejati, Crystal Lim, Dahlia Matouba, Atharva Inamdar, Christina Li, Hossein Khiabanian, Binfeng Lu, Ansu O. Perekatt

## Abstract

Intestinal tumorigenesis can occur via two distinct routes: *bottom-up* tumorigenesis occurs from mutations sustained in the Lgr5⁺ stem cells, whereas *top-down* tumorigenesis is driven by dedifferentiation of epithelial cells near the intestinal lumen. While sporadic human colon adenomas exhibit features of top-down tumorigenesis, their biological determinants remain elusive. Here, using a *Smad4* loss-of-function and *β-catenin* gain-of-function (Smad4^LOF^:β-catenin^GOF^) mouse model, we demonstrate that dedifferentiation-derived oncogenic stem cells sustain tumorigenesis more effectively than endogenous mutant stem cells harboring the mutation. The dedifferentiating villi epithelial cells showed early expression of CD44 and *Lgr5*, supporting oncogenic stemness. Aberrant Notch signaling in the villi epithelium was also detected at the onset of dedifferentiation, suggesting its contribution to dedifferentiation. Single-cell RNA sequencing revealed a distinct population of dedifferentiation-derived stem cells enriched for proliferative, metabolic, and mouse embryonic stem cell-like gene signatures, consistent with enhanced plasticity and tumorigenic potential. These mutant cells exhibited growth factor independence, indicating a capacity for niche-independent proliferation and metabolic adaptation to sustain tumor growth. These findings identify dedifferentiation-driven stemness, aberrant Notch activation, and metabolic plasticity as cooperative mechanisms that promote top-down intestinal tumorigenesis. This study provides insight into how oncogenic dedifferentiation contributes to tumor heterogeneity and persistence and has implications for therapeutic resistance in colorectal cancer.

## Introduction

Colorectal cancer (CRC) is one of the most prevalent and lethal malignancies worldwide, ranking second in global cancer-related mortality. Cancer stem cells (CSCs) are recognized as key drivers of tumor growth, metastasis, and therapy resistance, the latter contributing significantly to cancer relapse.^1–3^ Like normal intestinal stem cells, CSCs give rise to a hierarchical organization of the tumor tissue and are responsible for sustaining tumor growth.^4^

Lgr5⁺ stem cells are fast-cycling cells responsible for intestinal epithelial renewal. The intestinal epithelium is spatially organized, with the Lgr5⁺ stem cells and their proliferating progeny confined to the crypts (invaginations), while the villi (epithelial projections) house only the differentiated cells. Upon reaching the crypt–villus junction, progeny of Lgr5⁺ cells cease proliferation and differentiate. Despite this organization, the intestinal epithelium is highly plastic—differentiated cells can reacquire stemness during regeneration or upon Lgr5⁺ cell loss^5,6^. Similar fate plasticity occurs in colon tumors, where lineage-committed cells can regain Lgr5⁺ stem cell identity, highlighting niche occupancy as a key determinant of stem cell fate^7,8^.

The pathogenesis of CRC involves several key signaling pathways, with transforming growth factor-β (TGF-β) and Wnt cascades playing particularly critical roles.^2,9^ Smad4, a transcriptional effector of the Tgf-β and Bmp signaling pathway, suppresses growth, while β-catenin is the transcriptional effector of Wnt signaling, which promotes growth. These pathways are essential for maintaining intestinal homeostasis and regulating stem cell behavior; consequently, their dysregulation represents a hallmark of colorectal tumorigenesis.^10^

Experimental mouse models support two prevailing models for the cell-of-origin in colon cancer. The bottom-up model posits that oncogenic mutations in stem cells initiate tumorigenesis.^11^ Conversely, the top-down model suggests that dedifferentiation from the lineage committed or mature cells harboring mutations leads to tumor formation.^12,13^ The Verzi group previously demonstrated that simultaneous induction of Smad4 loss-of-function and β-catenin-gain-of-function (Smad4^LOF^:β-catenin^GOF^) mutations induces dedifferentiation, promoting proliferation and stemness within the villus compartment and ultimately leading to tumorigenesis.^14^ Given that the histological features of sporadic human colorectal tumors suggest a luminal origin, aligning with the top-down model, we used the Smad4^LOF^:β-catenin^GOF^ mouse model to investigate the factors favoring dedifferentiation-driven tumorigenesis and its implications. ^15^

scRNA-seq of dedifferentiating villus cells revealed heterogeneous stem-like populations encompassing proliferative and quiescent states. Moreover, we identified villus-specific alterations favored dedifferentiation-driven luminal tumorigenesis. Stem-cell–specific induction of the Smad4^LOF^:β-catenin^GOF^ showed a competitive disadvantage of mutant crypt stem cells, therby favoring luminal tumorigenesis via dedifferentiation.

Our findings provide insights into the mechanisms sustaining luminal tumorigenesis and highlight the potential role of dedifferentiation-driven stem cell heterogeneity in sustaining tumor growth.

## Methods

### Mouse model

All animal experiments were conducted according to protocols approved by the Institutional Animal Care and Use Committees (IACUC) of Stevens Institute of Technology. Mice were maintained under standard housing conditions with a 12-hour light/dark cycle, and all tissue collections were performed around midday to prevent diurnal variations in gene expression. To create the Smad4 loss-of-function: β-catenin gain-of-function (Smad4^LOF^:β-catenin^GOF^) conditional mutant model, the Villin-CreERT2 transgene^16^ was integrated into *Smad4^fl/fl^* ^17^ and *Ctnnb^exon3fl/wt^* conditional-mutant mice with a C57BL6-enriched background.^18^ Splicing out the floxed (fl) Smad4 allele and the exon 3 of the β-catenin allele is induced *via* tamoxifen injection. For Lgr5+ stem cell-specific induction of the double mutation, *Lgr5-EGFP-ires-CreERT2* allele was integrated into the *Smad4*^fl/fl^:*Ctnnb ^exon3fl/wt^* mouse.^19^

Two tamoxifen protocols were used based on experimental goals. For pan-epithelial mutation induction, tamoxifen was administered intraperitoneally at 0.05 g/kg for four consecutive days. For long-term mosaic induction, a single 0.01 g/kg injection produced palpable tumors within 1–2 months. Uninjected littermates served as wild-type controls. Both sexes were used, with age- and sex-matched replicates across groups.

between the wild type and the double mutant comparisons.

### Whole Intestinal Tissue Collection and Histological Processing

For histology and immunohistochemistry, freshly isolated small intestines were flushed with PBS, opened longitudinally, rolled, and fixed overnight in 4% paraformaldehyde at 4 °C. Tissues were dehydrated, processed in xylene, embedded in paraffin, and sectioned at 5 µm for analysis.

### Isolation of Intestinal Epithelium

For bulk RNA extraction and scRNA-seq, freshly harvested duodena were flushed with PBS, opened, and cut into ∼1 cm pieces. Epithelia were separated from mesenchyme by chelation in 5 mM EDTA/PBS for 20 min at 4 °C with rotation, followed by vigorous shaking. Villi were collected on a 70 µm filter, washed with ice-cold PBS, and pelleted at 200 rcf for 2 min at 4 °C. Crypts, obtained from the flow-through, were similarly washed and pelleted.

### Single-Cell Isolation

The villi and/or crypts collected as above, were digested in 2 ml dispase containing 200 µl DNase at 37 °C for 30 min with rotation. The suspension was shaken to release single cells, filtered through a 40 µm strainer, and pelleted at 300 rcf for 5 min at 4 °C. Single-cell processing followed according to downstream applications.

### Immunostaining Protocols

Immunohistochemistry was performed on 5 µm paraffin sections as described previously^14^. Primary antibodies against Smad4, CD44, EphB2, Glutaminase, TOM20, NICD, Lysozyme, and Cdx2 were used at dilutions listed in Tables ST1–ST2. Hematoxylin or methyl green served as nuclear counterstains. Fluorophore-conjugated secondary antibodies were applied for immunofluorescent staining, and nuclei were counterstained with DAPI.

### Hypoxia Detection

To visualize tissue hypoxia, mice were injected with Hypoxyprobe (0.1 mg/g body weight) one hour before sacrifice. The tissues were paraffin-embedded and sections were processed for fluorescent immunohistochemistry to detect Hypoxyprobe adducts.

### Image Acquisition and Processing

Brightfield images were captured using a Nikon Eclipse Ci-L microscope with a DS-Fi3 camera. Fluorescent images were acquired on a Zeiss LSM 880 confocal microscope under consistent laser and imaging settings. Brightness and contrast adjustments, when applied, were uniformly implemented across comparable samples.

### Western Blotting

The isolated villi epithelium were lysed in RIPA buffer (20 mM HEPES, 150 mM NaCl, 1 mM EGTA, 1% Triton X-100, 1 mM EDTA) supplemented with protease inhibitors (1× PI, 20 mM NaF, 1 mM Na₃VO₃, 1 mM PMSF). Lysates were rotated for 30 min at 4 °C and centrifuged at 21,130 rcf for 15 min to obtain soluble protein. Protein concentration was measured by BCA assay, and antibodies with corresponding dilutions are listed in Tables ST3–ST4.

### Organoid Culture

Organoids were established from villi and crypts of double mutant (DM) mice, with wild-type crypt organoids as controls. Villi-derived organoids were generated as described previously^20^. For crypt-derived organoids, freshly isolated duodena were flushed, opened, cut into ∼1 cm pieces, and incubated in 3 mM EDTA/PBS for 50 min at 4 °C with rotation. Crypts were released by shaking, filtered through a 70 µm strainer, washed, and pelleted at 200 rcf for 2 min at 4 °C. Crypts were embedded in Matrigel (10 crypts/µl) and cultured in either complete medium or medium lacking EGF, Noggin, and R-spondin. Reagents used are listed in Table ST6.

### Bulk RNA Sequencing

For bulk RNA sequencing, villus epithelium was collected and processed for RNA extraction using TRIzol. Library preparation, sequencing, and quality control were performed by Novogene (Sacramento, CA). Samples with RIN >4 were used for library construction and sequenced on an Illumina NovaSeq platform to generate 150 bp paired-end reads. Clean reads were processed using Novogene’s in-house Pearl pipeline, aligned to the *mm9* genome with HISAT2 v2.0.5, and quantified using featureCounts v1.5.0-p3.

### Single-Cell RNA Sequencing

Single cells prepared as described were washed in 0.1% BSA/PBS and processed with the Miltenyi Biotec Dead Cell Removal Kit. Live cells were pelleted, washed in 0.5% BSA/PBS, and stained with CD44-FITC (1:20) on ice for 20 min, followed by DAPI (1:25,000) counterstaining. RFP⁺ proliferating and CD44⁺ stem-like cells were isolated by FACS and submitted for scRNA-seq at the Rutgers Genomics Center and the Molecular and Genomics Informatics Core, NJMS. Single-cell capture used the 10x Genomics 3’v3 platform, sequenced on the Illumina NovaSeq 6000. Barcode deconvolution and alignment were performed using Cell Ranger v6.0.1.

### Bioinformatics

#### Bulk RNA-Seq Analysis

Differential expression analysis was conducted in R (v4.3.1) using DESeq2 (v1.40.2)^21,22^. Genes with adjusted *p* < 0.05 (Benjamini–Hochberg correction)^23^ were deemed significant. Normalized counts from DESeq2 (‘counts(dds, normalized=TRUE)’), were used for Gene Set Enrichment Analysis (GSEA) via the Broad Institute GSEA-P application (v4.3.2)^24^. Mouse Ensembl IDs were mapped to human orthologs (MSigDB v2023.1.Hs), with 10,000 gene set permutations and default size filters (15–500 genes). Gene sets with FDR *q* < 0.05 were considered significantly enriched. Gene sets were sourced from the Molecular Signatures Database (MSigDB).^25^

#### Single-Cell RNA-Seq Data Analysis

Single-cell analyses were performed in Python using Scanpy (v1.11.1) ^26^. the epithelial cell types were annotated based on curated canonical marker gene sets, and gene set scores were computed with score_genes. Scores were visualized on UMAPs and summarized by dot plots. Differential expression between clusters was determined using the Wilcoxon rank-sum test with Benjamini–Hochberg correction (*adj. p* < 0.05). Pre-ranked GSEA based on Wilcoxon statistics was conducted using MSigDB and custom literature-derived gene sets.

### Materials

All materials used in this study are summarized in supplementary tables ST1-ST7.

## Results

### Dedifferentiation-induced acquisition of stemness in the villus epithelium promotes luminal tumorigenesis

Reacquisition of stemness is a hallmark of malignant transformation^27^. The Smad4^LOF^: β-catenin^GOF^ (double mutant) was induced throughout the intestinal epithelium using Villin-Cre to assess dedifferentiation-driven oncogenic stemness. Within one week, CD44, a putative cancer stem cell marker^28^, was detected in the villus epithelium, indicating oncogenic stemness (Fig. 1A). Because pan-epithelial induction (0.05 g/kg Tamoxifen for 4 days) was lethal within two weeks, a single low dose (0.01 g/kg) was used to generate mosaic mutants. Palpable tumors appeared within 1–2 months (Fig. 1B). Hyperplastic luminal regions displayed EphB2⁺ ectopic crypts (Fig. 1C, D), and the prevalence of luminal hyperplasia with invaginations indicated dedifferentiation near the lumen. The expression of CD44 and EphB2, both colon cancer stem cell markers^29,30^, suggests dedifferentiation-driven tumorigenesis via reacquisition of stemness.

**Figure 1.**
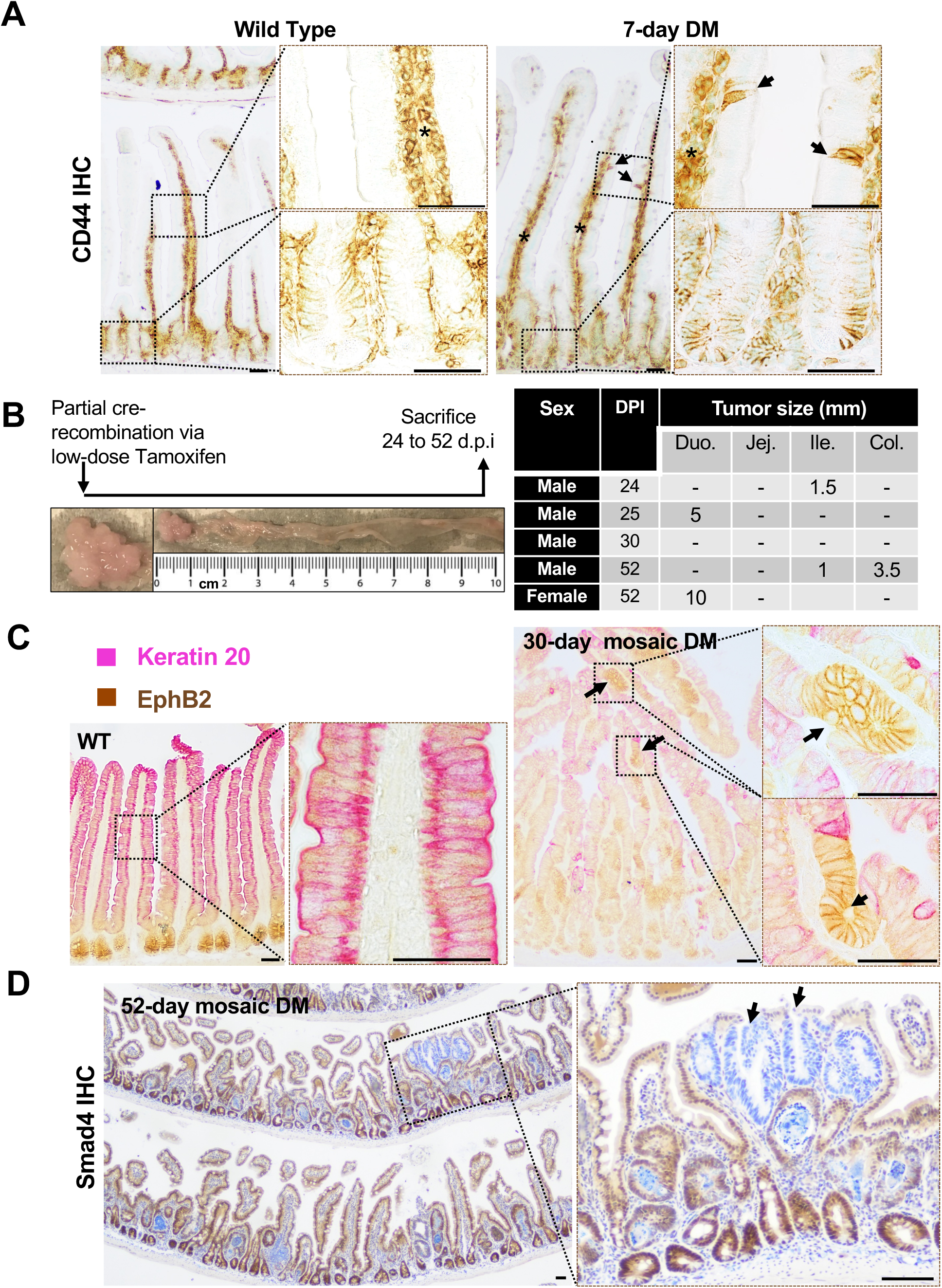
Dedifferentiation-driven tumorigenesis in the Smad4^LOF^: β-catenin^GOF^ double mutant. **A**, CD44 immunoreactivity in the double mutant villi epithelium (arrows), suggesting oncogenic stemness in the 7-day double mutant (DM) villi n = 3. **B,** left panel: Schema for mosaic induction of the double mutation (DM), and representative image of the resulting adenoma. Right panel: region within the intestine where tumors were found and their size. **C,** Co-immunostaining for Keratin 20 (pink) and EphB2 (brown) in the wild type (left) and the mutant (right). Arrows point to the nascent ectopic crypts within the tumor. **D,** IHC showing loss of Smad4 in the hyperplastic regions near the lumen (arrows). Boxed regions within the image are enlarged to the right. * Non-specific CD44 immunoreactivity in the mesenchyme. Scale bars, 50 μm.

### Transcriptional shifts in the dedifferentiating double mutant villi epithelium support oncogenic stemness

To define epithelial transcriptional changes driving dedifferentiation and oncogenic stemness, we compared the transcriptomes of double mutant and wild-type villus epithelia at 7 days post-induction, when CD44⁺ cells and nascent ectopic crypts first appear (Fig. 1A). Downregulation of the tumor suppressor *Clca4a* and upregulation of the stem cell marker *Prom1*^31,32^, along with altered expression of β-catenin and Smad4 targets, confirmed successful induction of the β-catenin^GOF^: Smad4^LOF^ mutation (Fig. 2A, B). RNA-seq and Gene Set Enrichment Analysis (GSEA) revealed enrichment of Hallmark Myc and E2F pathways^25^, as well as gene signatures associated with cell fate plasticity and tumor-related embryonic stemness^31,32^ (Fig. 2B). These results align with the established roles of E2F and Myc signaling in driving cellular reprogramming and stemness during tumorigenesis. The enrichment of cell fate plasticity and embryonic stemness signatures indicates mutation-induced transcriptional reprogramming that promotes oncogenic stemness in the dedifferentiating villus epithelium^35–38^.

**Figure 2.**
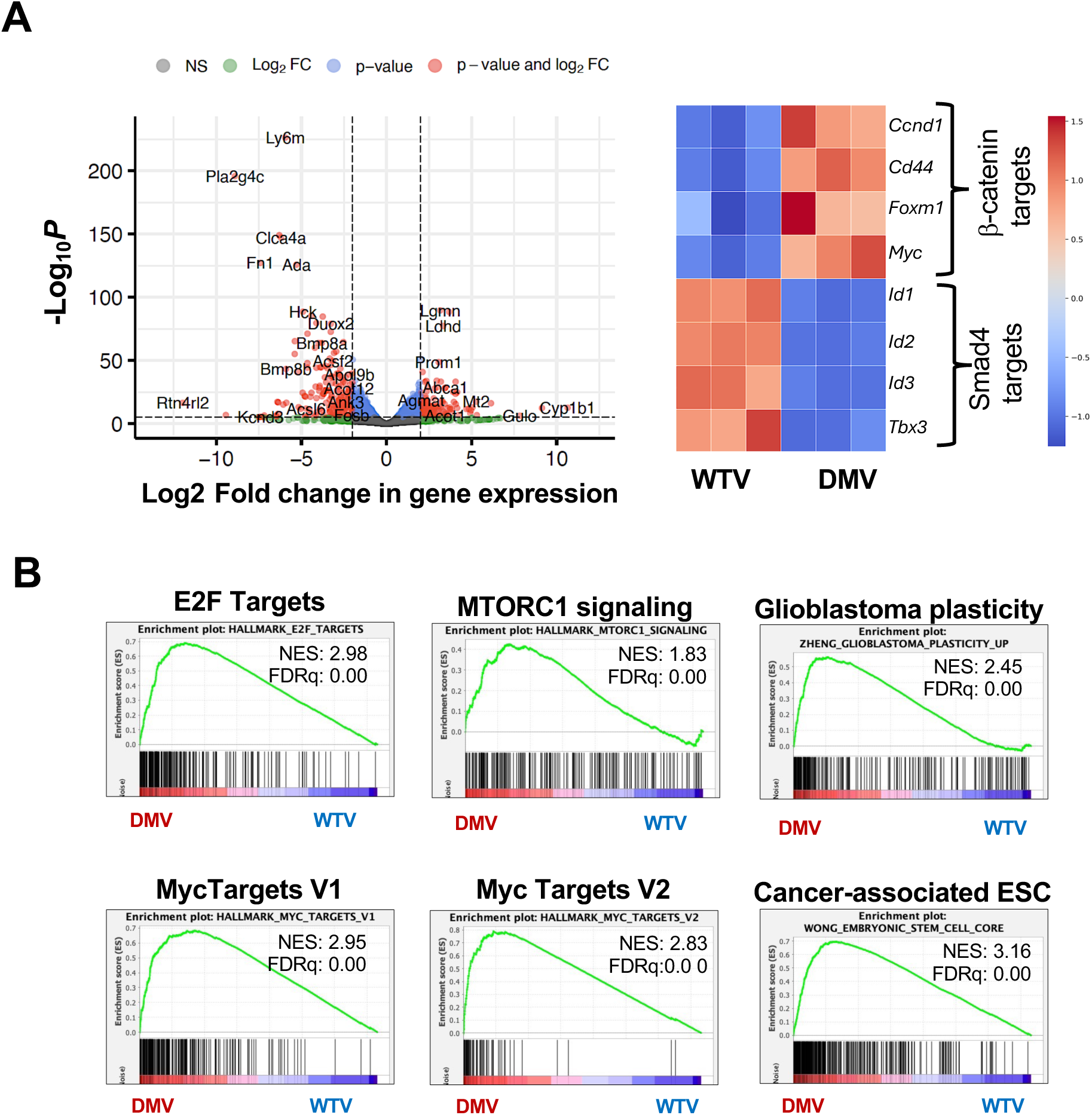
Early transcriptional alterations support oncogenic stemness and tumorigenesis in the villi epithelium within seven days after inducing the Smad4^LOF^: β-catenin^GOF^ mutation. **A**, Volcano plot of differentially expressed genes in the villi epithelium seven days after the mutation. **B,** Heatmap confirming differential expression of Smad4 and β-catenin transcriptional targets in the double mutant villi (DMV) vs wild type villi (WTV) epithelium, validating simultaneous mutation induction. **C**, GSEA plots showing enrichment of the Hallmark Myc and E2F targets, mTORC1 signaling, and gene signatures of stemness and plasticity, consistent with the induction of tumorigenic potential in the mutant villi. NES = normalized enrichment score, FDRq = False discovery rate q-value.

### The Lgr5 stem cell drift dynamics are skewed against the persistence of the double stem cells in the endogenous crypts

Lgr5⁺ stem cells undergo neutral drift dynamics in the normal intestinal epithelium, dividing symmetrically to replace each other and maintain clonal crypts ^39,40^ To assess how the Smad4^LOF^: β-catenin^GOF^ mutation affects tumorigenesis when induced specifically in Lgr5⁺ stem cells, we used the Tamoxifen-inducible Lgr5-Cre^ERT2^;EGFP mice^19^, which enable visualization of the subset of Cre-activable Lgr5^+^ stem cells by GFP immunostaining. At 10 days post-induction (d.p.i.), Smad4⁻ cells appeared within crypts, confirming successful mutation induction (Fig. 3A). By 22 d.p.i., these double mutant cells in the crypts were largely replaced by wild-type cells, indicating a competitive advantage for the wild type Lgr5^+^ stem cells in the crypts. Notably, CD44 and Lgr5⁺ cells, evidenced by GFP⁺ immunoreactivity, appeared in the Smad4⁻ hyperplastic regions near the lumen(Fig. 3B), suggesting oncogenic stemness and adenoma initiation. This contrasting distribution—loss of mutant stem cells from the endogenous crypts but their appearance near the lumen—indicates that the villus region provides a more favorable niche for tumor growth.

**Figure 3.**
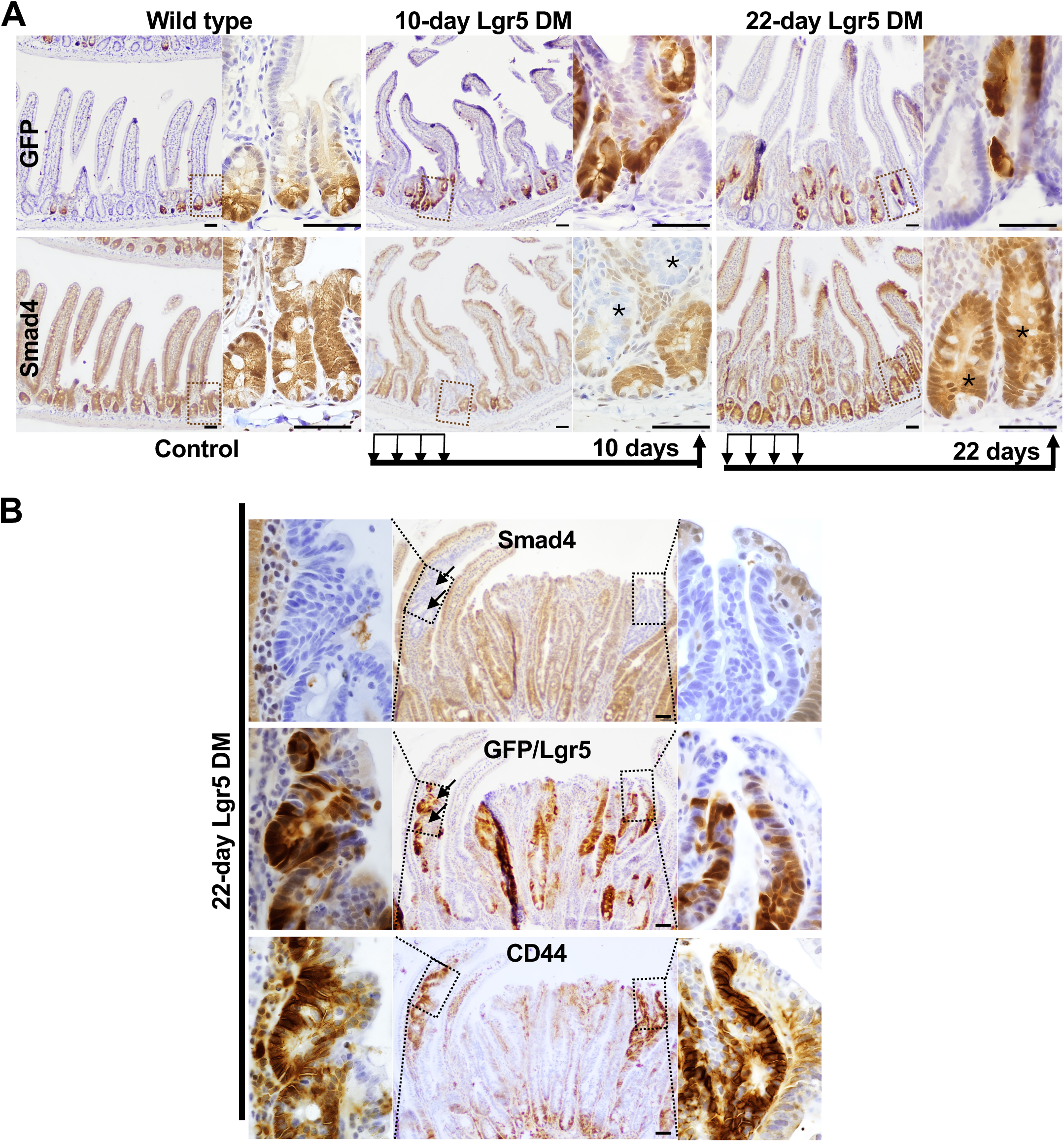
Loss of mutant Lgr5+ stem cells from the endogenous crypts and dedifferentiation-driven Lgr5+ stem cells near the lumen. **A**, GFP (top) and Smad4 (bottom) IHC showing Lgr5^+^ and wild type stem cells on adjacent sections before and after Lgr5^+^ stem cell-specific induction of the mutation; the Lgr5-driven Cre in *Smad4f^l/fl^; Ctnnb^exon3fl/wt^; Lgr5-EGFP-ires-CreERT2* mice enabled the mutation only in a subset of the Lgr5^+^ stem cells and stem cell visualization via GFP expression. Note the presence of mutant cells (asterisk) at 10 days but not 22 days in the endogenous crypts. **B,** Smad4 (top), GFP (middle) and CD44 (bottom) immunostaining on adjacent sections, showing Lgr5⁺ stem cells in the Smad4-deficient hyperplastic regions near the lumen. Note the GFP expression in the villus invaginations (arrows), suggesting the emergence of Lgr5+ stem cells via dedifferentiation. Boxed regions are enlarged to their immediate left and right. n = 3 per time-point. Scale bar = 50 μm

### The mutation triggers metabolic alterations specifically in the villi that facilitate tumor growth

To identify factors favoring luminal tumorigenesis, we first assessed hypoxia, which increases along the crypt–villus axis in the normal epithelium^41^. Pimonidazole labeling revealed elevated hypoxia in the double mutant villi but not in crypts or ectopic crypts, indicating a transient, pre-ectopic crypt event (Fig. 4A). Because glutamine metabolism can induce hypoxia through increased respiration^42^, we probed for glutaminolysis and mitochondrial function. The double mutant villi showed increased Glutaminase (GLS) and Tom20 expression (Fig. 4B, C), indicating elevated glutaminolysis and mitochondrial function^43,44^. Consistent with enhanced oxidative metabolism-associated cytoprotective activity, the antioxidants Prdx3 and Prdx6 and the mitochondrial quality-control protein PINK1 were upregulated (Fig. 4D) ^42,45–47^. These findings suggest that metabolic reprogramming and associated cytoprotective mechanisms in the villus epithelium promote dedifferentiation-driven tumorigenesis near the lumen.

**Figure 4.**
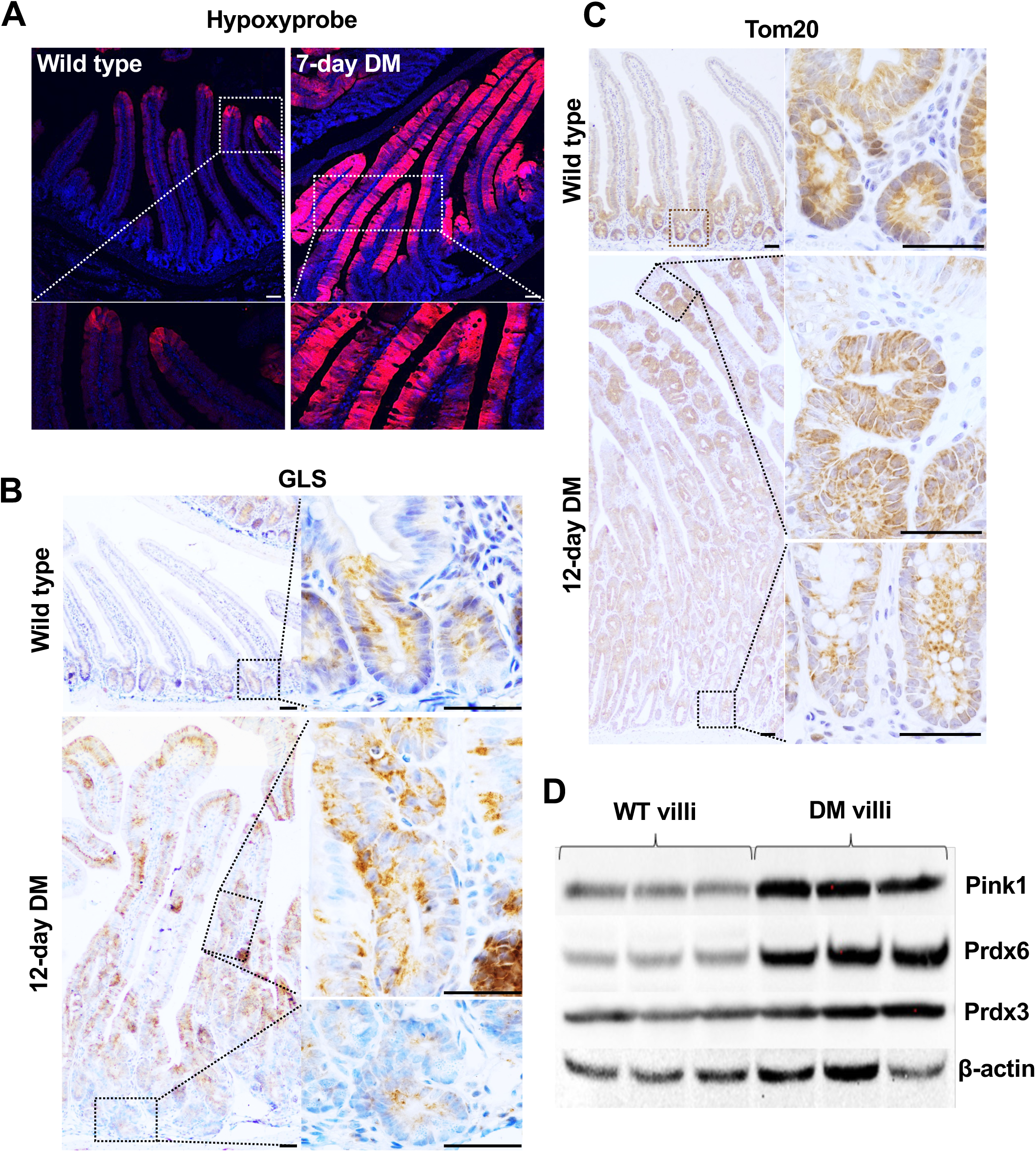
Mutant villi-specific metabolic changes that support tumor growth. **A,** Pimonidazole adduct immunostaining showing increased hypoxia in the double mutant villi. **B,** Elevated glutaminase in mutant villi but not the crypts, indicated by GLS immunoreactivity. **C,** increased TOM20 immunoreactivity, indicative of increased mitochondrial biogenesis. **D,** Immunoblotting for the glutamine metabolism-associated antioxidants, PRDX3 and PRDX6, and the mitochondrial sentinel, Pink1 in the villi epithelial lysates, with β-actin as the loading control. N = 3 per treatment. Scale bars, 50 μm. DM = tamoxifen injected *Smad4f^l/fl;^ Ctnnb^exon3fl/wt^*; *Villin^Cre^ERT2*.

### Aberrant Notch signaling in the double mutant villi precedes ectopic crypt formation

In the normal intestinal epithelium, Notch signaling is confined to the crypts and activated by ligands expressed on adjacent Paneth cells^48^. Because Notch activity is essential for intestinal stem cell maintenance, we examined the Notch intracellular domain (NICD) as a marker of active signaling^49^. As expected, NICD immunoreactivity was observed in the endogenous crypts. Unexpectedly, the double mutant villus epithelium also showed NICD staining within seven days of mutation induction—before ectopic crypt formation (Fig. 5). Immunostaining for the Paneth cell marker lysozyme on adjacent sections revealed Paneth cells in the mutant crypts but not in the mutant villi at this stage. Thus, the presence of NICD in the absence of Paneth cells in the villi indicates aberrant, Paneth cell–independent Notch activation at the onset of dedifferentiation, implicating Notch signaling in promoting stemness in the double mutant villus epithelium.

**Figure 5.**
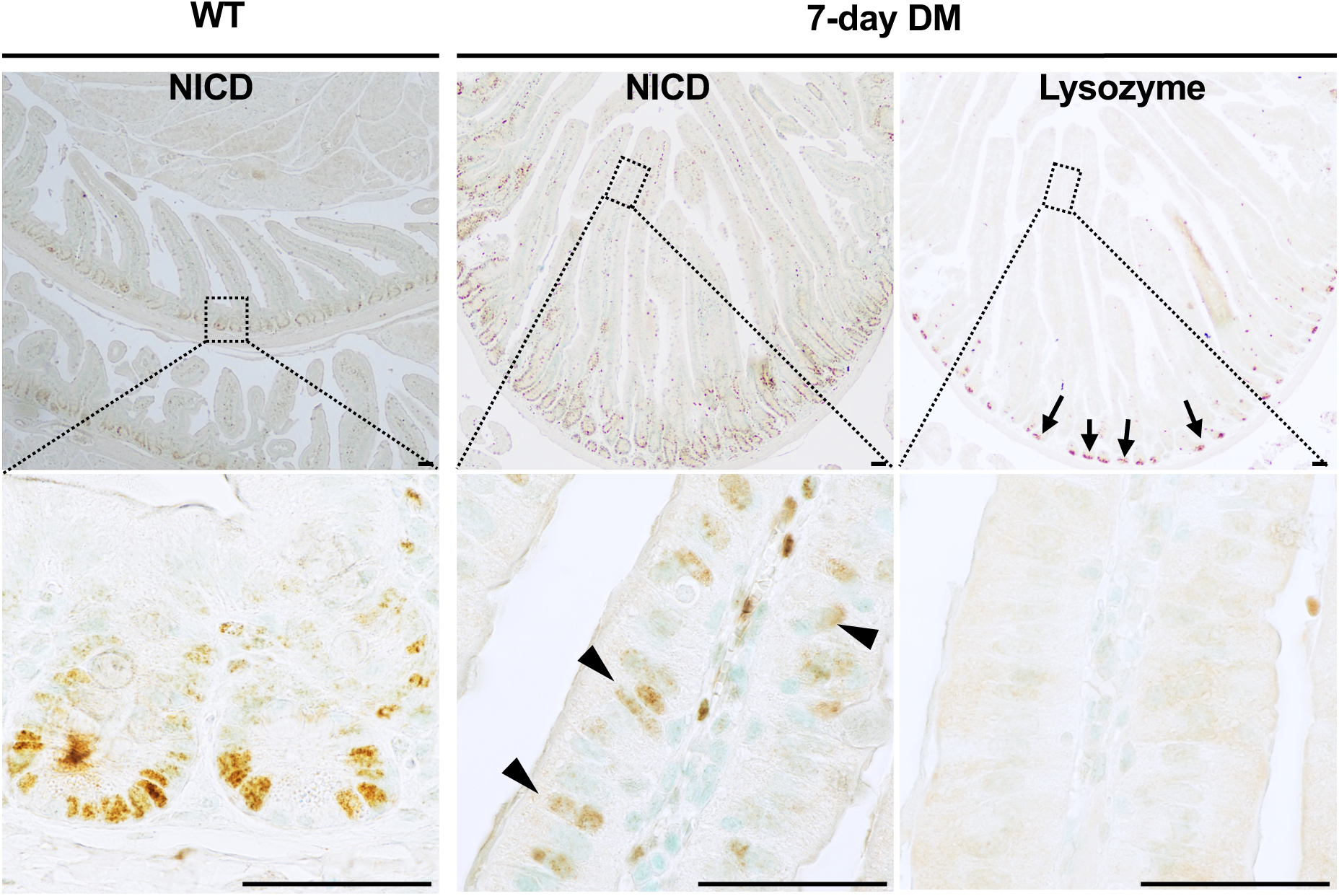
Paneth cell-independent Notch signaling in the mutant villi at the onset of dedifferentiation. Left panel: Notch signaling, indicated by NICD immunoreactivity in the wild type (WT) crypts. The middle and right panels show NICD and Lysozyme immunostaining, respectively, on adjacent tissue sections of the 7-day double mutant intestine. Notch signaling is apparent in the villi (arrowheads) within seven days of inducing the mutation, but Paneth cells, evidenced by Lysozyme immunoreactivity (arrows), are restricted to the crypts. The boxed regions in the top panel are magnified in the bottom panel. n = 3 per treatment. DM = tamoxifen injected *Smad4f^l/fl;^ Ctnnb^exon3fl/wt^; Villin^Cre^ERT2* mice. Scale bars, 50 μm.

### Dedifferentiation in the double mutant epithelium leads to stem-like cells with distinct transcriptional profiles

Despite pan-epithelial induction of the double mutation, proliferative or stem cell marker expression in the villus epithelium was sporadic. To enrich these populations for single-cell RNA sequencing (scRNA-seq), proliferating cells were genetically labeled with RFP^50^, and seven days post-induction, crypt and villus epithelia were dissociated and immunolabeled with GFP-conjugated CD44. RFP⁺ and/or GFP⁺ cells were isolated by FACS and subjected to scRNA-seq (Fig. 6A). Control samples comprised RFP⁺ and/or CD44⁺ crypt cells and an equal number of villus epithelial cells from wild-type mice. Leiden clustering identified 15 populations (10,403 cells) in wild type, 13 (4,143 cells) in mutant villi, and 12 (7,860 cells) in mutant crypts (Fig. 6B, S2–S3). Based on composite expression of the Lgr5+ stem cell markers^19,51^ *Gkn3*, *Lgr5*, and *CD44* (Stem_score), clusters 5 and 7 were designated stem populations in wild type, while the mutant villi contained five (c3, c6, c9, c10, c14) and the mutant crypts four (c3, c6, c9, c10) stem clusters, indicating greater stem cell heterogeneity in the mutant epithelium (Fig. 6C, S2A, S3B & C).

**Figure 6.**
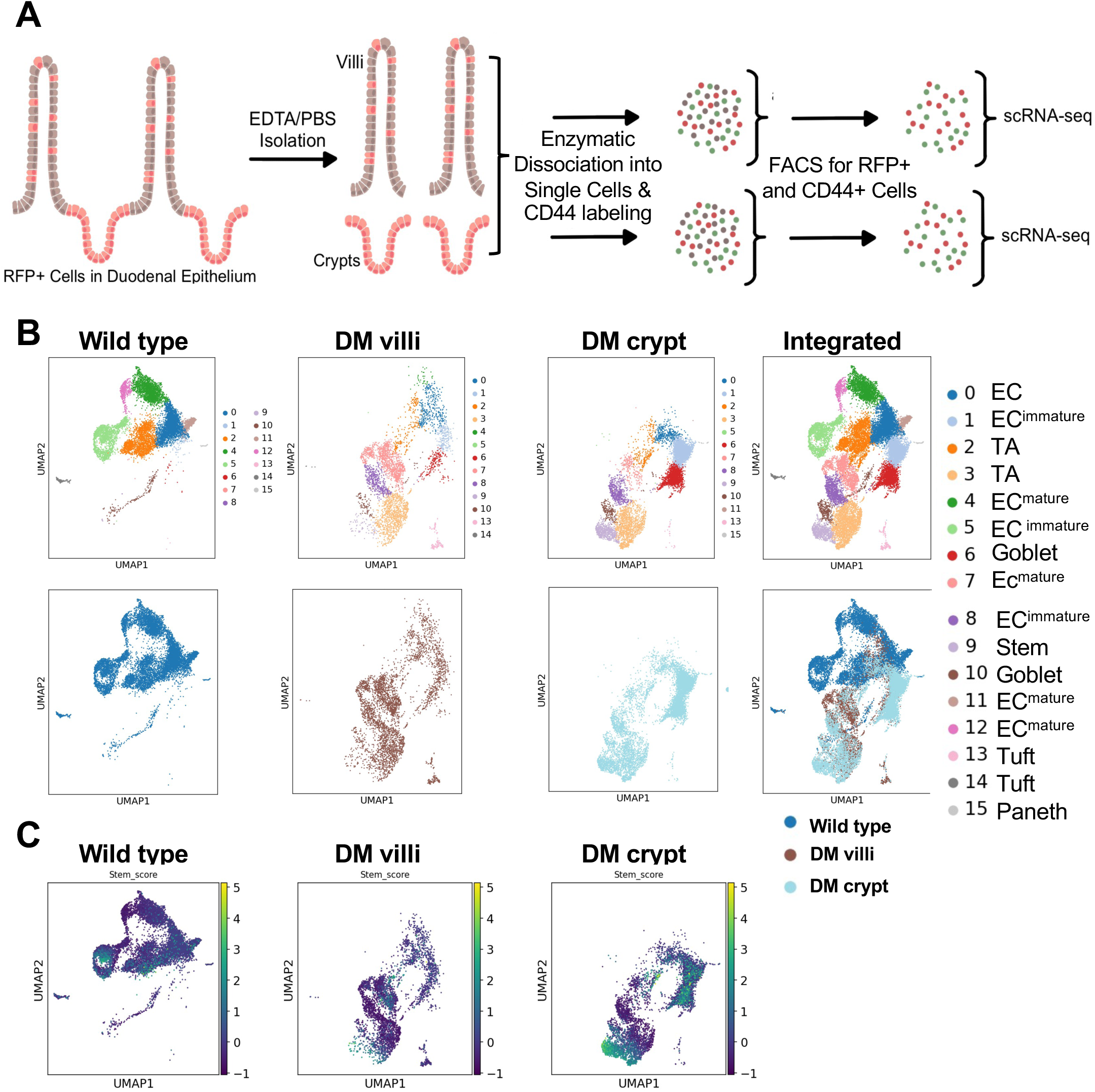
Dedifferentiation in the mutant epithelium leads to stem-like cells with distinct transcriptional profiles. **A**, Workflow of sample preparation and processing for scRNA-seq. **B**, Leiden clustering and batch integration, revealing populations with shared and distinct transcriptional profiles in the wild type (crypt and villi) and the dedifferentiating cells in the double mutant (DM) villi and crypt epithelium. **C**, Stem cell clusters, indicated by the composite expression levels of the Lgr5^+^ stem cell markers Gkn3, Lgr5, and CD44 (Stem_score) in the three samples. DM = tamoxifen injected *Smad4f^l/fl;^ Ctnnb^exon3fl/wt^; Villin^Cre^ERT2* mouse.

### The heterogeneity among the stem populations reflects differences in proliferation, metabolic state, and signaling

Stem cell heterogeneity enhances adaptability and growth potential. As the transcriptional profiles of double-mutant stem clusters differed from the wild type, we compared them to identify features of dedifferentiation-derived stemness. Embryonic stem cell (ESC)-like gene signatures, reported in human cancers^34^, were evaluated across double mutant villus stem clusters (c3, c6, c9, c10, c14) relative to wild-type cluster c5 (1,866 cells; Fig. S2B). Only cluster c3 showed significant enrichment for the cancer-associated ESC module (Fig. 7A). Consistent with the mESC-like signature, the mutant villi epithelium exhibited heterogeneous loss of the intestinal lineage factor Cdx2 (Fig. 7B). Cluster c3 was also enriched for Hallmark E2F, MYC, and MTORC1 signaling, reflecting higher proliferative and metabolic activity in c3 (Fig. 7C). In contrast, clusters c9, c10, and c14 showed enrichment for Hedgehog signaling but not proliferative signatures, suggesting survival signaling^52^ in these clusters (Fig. 7D). Since c3 was unique to the double mutant epithelium, comparison of villus and crypt c3 populations revealed enrichment for reactive oxygen species and *mTORC1* pathways in the double-mutant villus c3, highlighting metabolic distinctions between the double-mutant villus and crypt epithelial stem populations (Fig. 7E).

**Figure 7.**
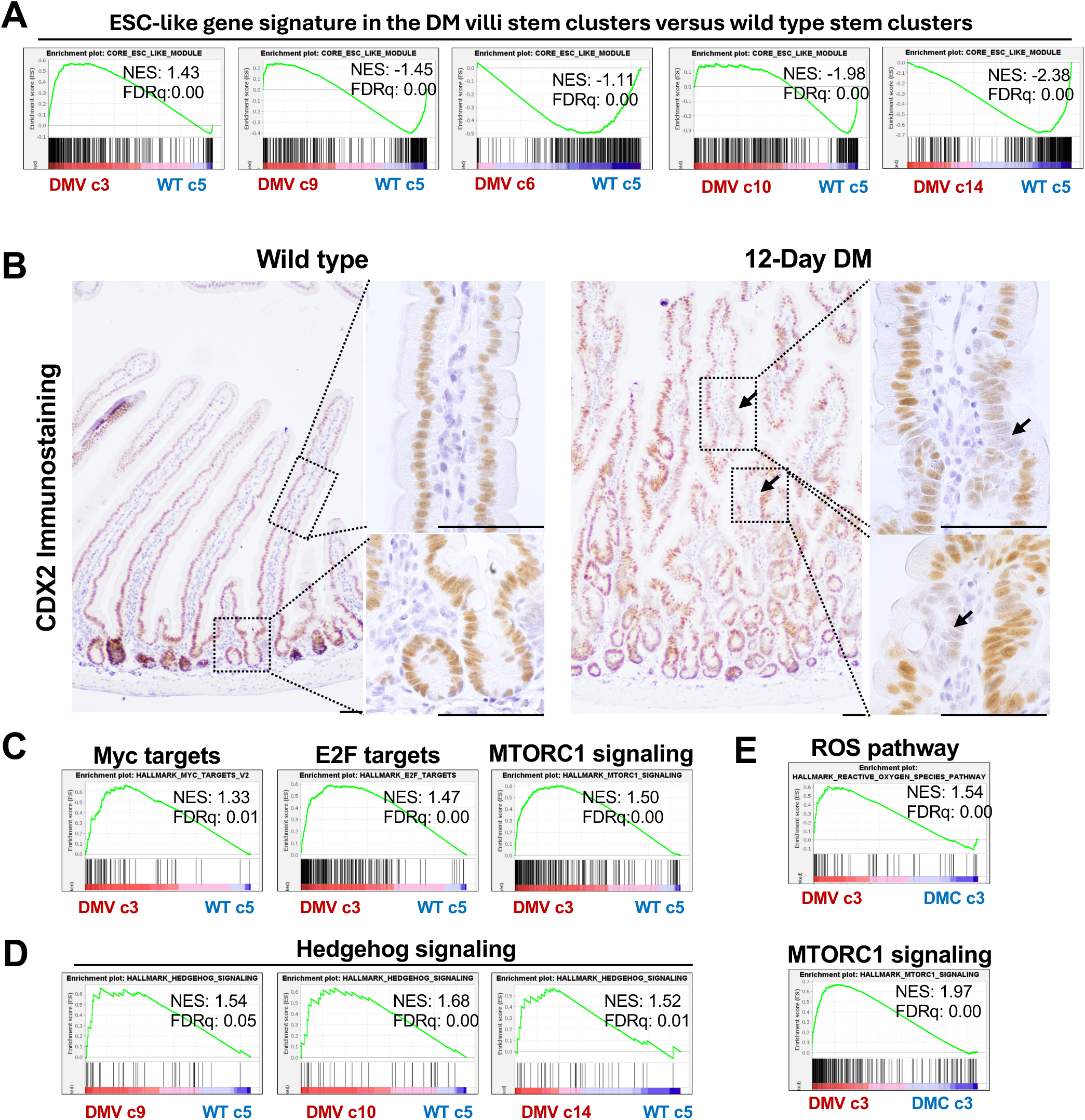
Differential enrichment of proliferative and metabolic gene signatures in the mutant stem populations. **A**, GSEA plots showing enrichment of Embryonic Stem Cell (ESC)-like program exclusively in the mutant villi stem cluster 3 (DMV c3) relative to the wild type stem cluster 5 (WT c5); note negative enrichment of the ESC-like gene signature in the rest of the double mutant villi stem clusters. **B**, sporadic loss of Cdx2, in the double mutant, indicative of the loss of the intestinal lineage specification in a subset of the mutant cells (arrows). **C**, GSEA plots showing enrichment of the proliferative and metabolic gene signatures in the double mutant villi stem cluster 3 (DMV c3) compared to the wild type stem cluster 5 (WT c5), and **D**, GSEA plots showing the enrichment of the Hedgehog signaling gene signature in mutant villi clusters, c9, c10, and c14 relative to the wild type stem cluster 5 (WT c5). Differences between the stem cluster 3 of the mutant villi and mutant crypts, evidenced by GSEA plot showing enrichment of the metabolic gene signatures in the double mutant villi stem cluster 3 relative to the double mutant crypt stem cluster 3 (DMV c3 vs DMC c3). NES = normalized enrichment score, FDRq = False discovery rate q-value.

## Discussion

The bottom-up and the top-down are the two models of intestinal tumorigenesis that differentiate the origin of tumors: the former, from mutations incurred in the stem cells, while the latter, owing to dedifferentiation. However, the sporadic colon tumors in human patients display the histological characteristics suggestive of dedifferentiation-driven tumorigenesis: the adenomatous polyps appear to develop from genetically altered cells in the superficial mucosae^15^. Several groups have reported top-down or luminal tumorigenesis in mouse models of intestinal tumorigenesis^12,53,47,48,49^. However, the implications of dedifferentiation-driven versus bottom-up tumorigenesis are unknown.

Using the Smad4^LOF^:β-catenin^GOF^ mouse model, we demonstrate that dedifferentiation-derived oncogenic stem cells more effectively sustain tumorigenesis. These cells appear to possess a competitive advantage over endogenous mutant stem cells carrying the same mutation. Given that Smad4^LOF^:β-catenin^GOF^ mutations mimics the effect of two of the most common colon cancer drivers in human patients, our mouse model provides insight into how tumors persist, driven by heterogeneity in proliferative and metabolic states, and distinct transcriptional states that confer the diversity to survive a changing environment.

CD44, a putative colorectal cancer stem cell marker implicated in adenoma initiation, ^56,57^ was sporadically expressed in the Smad4^LOF^:β-catenin^GOF^ mutant villus epithelium following pan-epithelial induction and before ectopic crypt formation (Fig. 1A), indicating villus-origin adenoma initiation. scRNA-seq further revealed Lgr5 expression in dedifferentiating villus cells (Fig. 6C, S3B), consistent with reports that non-Lgr5 cells can initiate tumors and regenerate Lgr5⁺ stem cells after depletion^8,56^.

Aberrant Notch signaling in the mutant villi indicates its involvement in dedifferentiation and adenoma initiation. Elevated Notch activity is reported in human adenomas and carcinomas^58,59^. Since Notch signaling in the normal intestine is confined to crypts and activated by Paneth cell ligands, its ectopic activation in villus epithelium (Fig. 5) before crypt formation and independent of Paneth cells suggests a role in triggering adenoma development, consistent with its requirement for generating self-renewing adenoma-initiating cells^60,61^. However, the source of the Notch-ligand and whether the Notch signaling is ligand-independent in the villi before ectopic crypt formation is unknown. Given that villus cells may express distinct NICD-associated transcription factors, identifying Notch-dependent transcriptional targets specific to tumorigenesis warrants further investigation.

Under normal conditions, stem cells undergo neutral drift dynamics, competing equipotently for niche occupancy to establish monoclonal crypts^40^. In contrast, the altered drift dynamics observed following Lgr5-Cre-driven Smad4^LOF^:β-catenin^GOF^ mutation (Fig. 3) indicate a competitive advantage of wild-type stem cells for niche occupancy. This was unexpected, as Apc loss and heightened Wnt signaling typically promote bottom-up tumorigenesis^11^. The likely cause is mutation-induced disruption of BMP signaling, which normally restrains stem cell proliferation^62^. Because Bmpr1 is expressed in stem cells but absent in their progeny, Smad4 loss is predicted to enhance proliferation, increasing the probability of mutant stem cells being displaced from the niche—an environment rich in EGF, Noggin, and R-spondin required for Lgr5⁺ stem cell maintenance^63^. The growth factor independence of mutant-derived organoids (Fig. S1) supports this notion, suggesting that mutant cells survive without niche signals. In contrast, wild-type cells rely on them and therefore remain niche-resident. Consequently, the highly proliferative, niche-displaced mutant cells are exposed to differentiation cues that promote lineage commitment^64^.

The mutation-induced metabolism-associated changes in the villi epithelium (Fig. 4) suggest that the villi-specific alterations favor tumor growth. Metabolic adaptations are essential for tumor growth to meet the high biosynthetic processes and energy demand in fast-growing cancer cells.^65,66^ Glutaminolysis also increases mitochondrial respiration, which might increase reactive oxygen species. Our data suggest increased mitochondrial function and the accompanying cytoprotective mechanisms in the villi epithelium, implicating metabolic vulnerabilities in top-down tumorigenesis.

scRNA-seq analysis revealed marked heterogeneity among stem cell populations, reflecting differences in proliferation, signaling, and metabolism. The Smad4^LOF^:β-catenin^GOF^ mutation generated a distinct stem cell cluster (c3) with no transcriptional overlap with wild-type epithelia. Cluster c3 was enriched for cancer-associated, mESC-like gene signatures consistent with heterogeneous loss of the intestinal lineage factor Cdx2 (Fig. 7B). Co-enrichment of proliferative and metabolic pathways in mutant villi cluster 3 indicates active tumor-driving potential. In contrast, clusters 9, 10, and 14 lacked proliferative signatures, suggesting a quiescent state.

Our findings suggest that metabolic adaptations and dedifferentiation-driven stem cell heterogeneity may enhance adaptability and confer a selective advantage driving top-down tumorigenesis.

## Data Availability

Bulk RNA-seq data [GSE263192] and single-cell RNA-seq data from the double mutant epithelium [GSE288393] and wild type epithelium [GSE288394] have been submitted to the National Center for Biotechnology Information Gene Expression Omnibus.

## Acknowledgment

A. Perekatt was supported by NIH grant K22ACTFCA218462. We thank Dr. Michael P. Verzi for discussions, Dr. Angela L. Tyner for manuscript review, and the Genomics Center and Molecular and Genomics Informatics Core at Rutgers New Jersey Medical School for single-cell RNA sequencing support.

