## SUPPLEMENTARY FIGURES for "Dedifferentiation-Driven Oncogenic Stemness Promotes Tumor-Sustaining Adaptability in the Intestinal Epithelium"

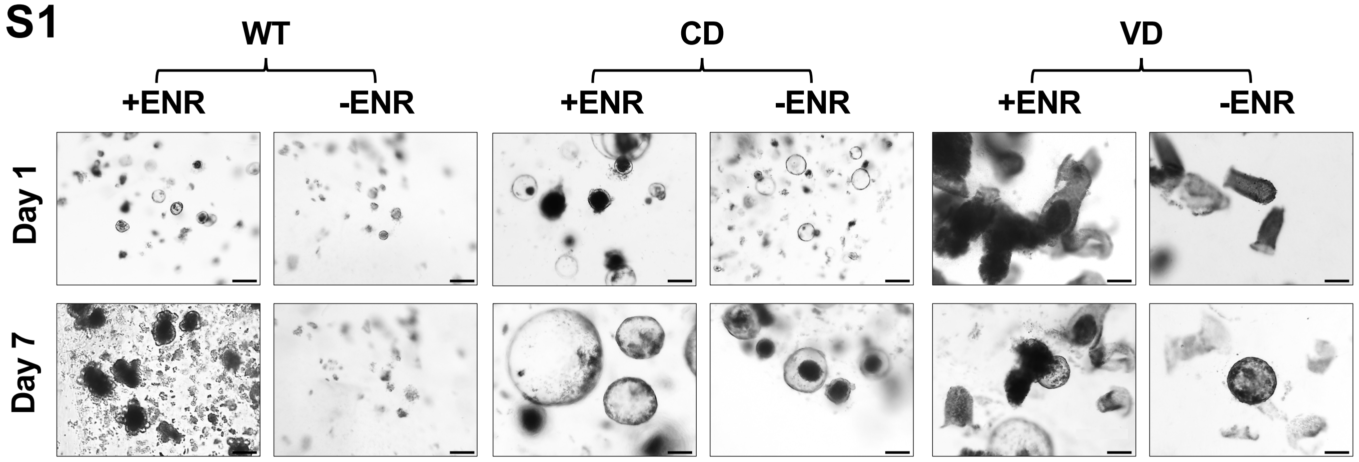


**Figure S1. Organoids from the double mutant villi epithelium can grow in the combined absence of Epidermal Growth Factor (E), R-spondin (R), and Noggin (N).** Wild-type crypt epithelium (WT), double-mutant crypt epithelium (CD), or the double-mutant villi (VD) were plated in the presence or the combined absence of the three growth factors (ENR). Unlike the wild type, which showed no organoids without the ENR, the crypt and villi-derived organoids from the mutant mouse showed growth factor independence seven days after plating.

A

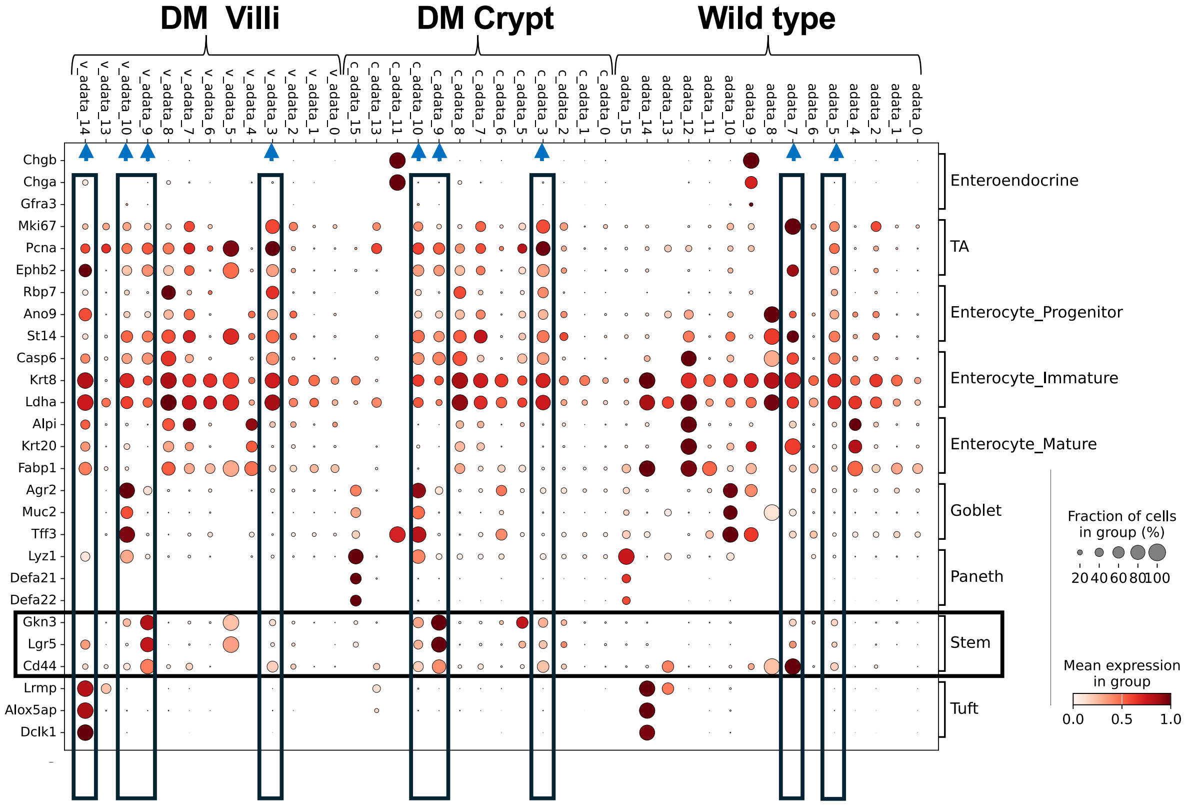


B

| c # | Wild type | DM villi | DM crypt |
|---|---|---|---|
| leiden | control_adata | v_adata | c_adata |
| 0 | 3193 | 375 | 388 |
| 1 | 72 | 216 | 2556 |
| 2 | 1987 | 231 | 261 |
| 3 | 0 | 1046 | 1287 |
| 4 | 2002 | 37 | 0 |
| 5 | 1866 | 1 | 6 |
| 6 | 17 | 207 | 1632 |
| 7 | 3 | 1229 | 227 |
| 8 | 1 | 480 | 536 |
| 9 | 7 | 76 | 688 |
| 10 | 172 | 106 | 242 |
| 11 | 497 | 0 | 1 |
| 12 | 410 | 0 | 0 |
| 13 | 3 | 135 | 23 |
| 14 | 153 | 4 | 0 |
| 15 | 20 | 0 | 13 |
| Total | 10403 | 4143 | 7860 |

**Figure S2. Data from batch integration analysis: A.** dot plot of various intestinal lineage markers expressed in the various clusters from each sample; blue arrows point to the clusters most enriched for Lgr5+ stem cell marker genes. The horizontal boxed region points to the Lgr5$^{+}$ marker gene expression. The size of each dot corresponds to the fraction of cells within a cluster expressing the indicated gene, while the color intensity represents the average expression level. and **B**, the relative number of cells in the various clusters of the three samples. control_adata = wild type epithelium, v_adata = mutant villi, c_adata = mutant crypt.

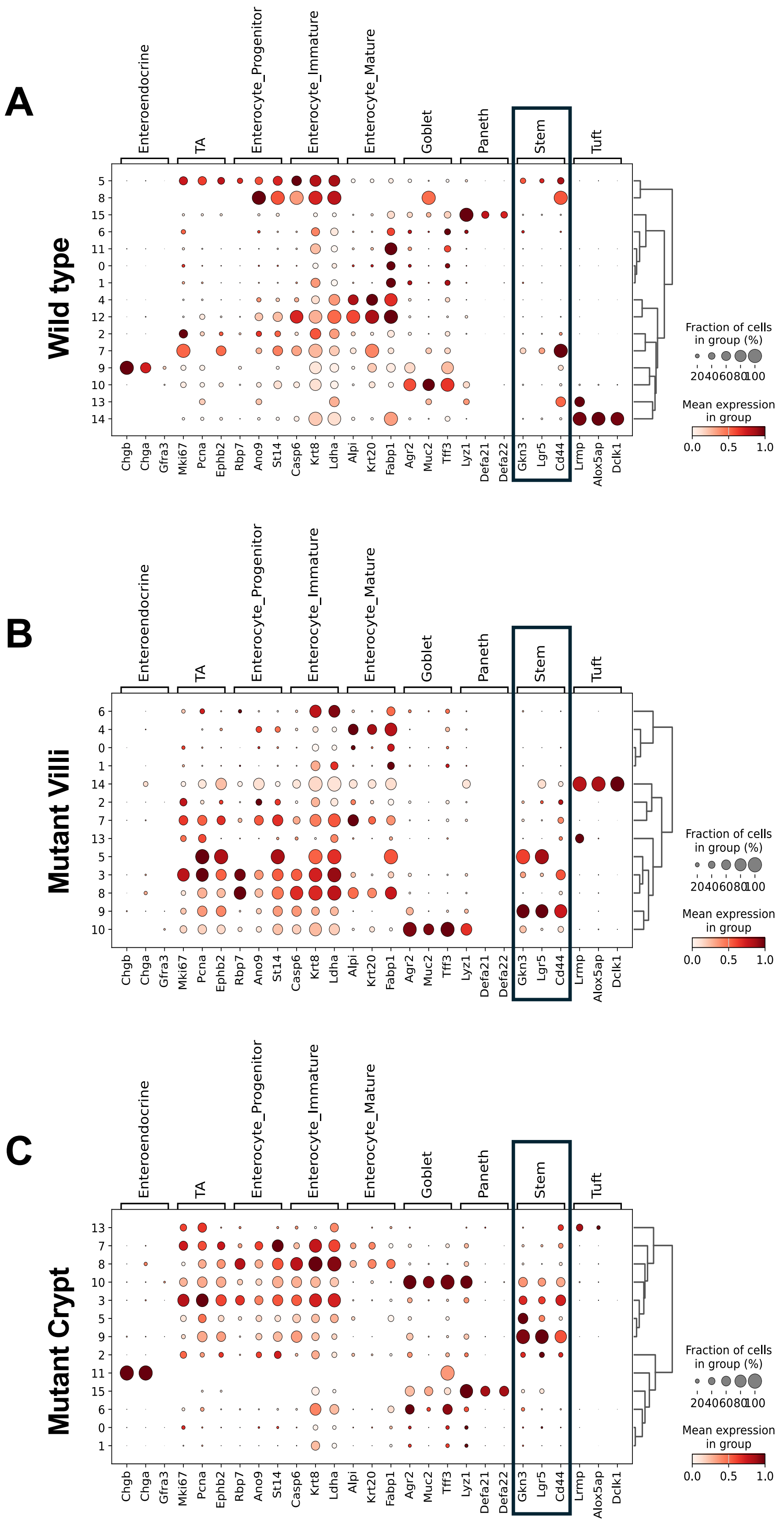


**Figure S3. Non-integrated analysis showing** dot plot of various intestinal lineage markers expressed in the various clusters of the wild type epithelium (**A**), dedifferentiating cells in the double mutant villi epithelium (**B**), and the double mutant crypts (**C**). The size of each dot corresponds to the fraction of cells within a cluster expressing the indicated gene, while the color intensity represents the average expression level.
