## SUPPLEMENTARY TABLES for "Dedifferentiation-Driven Oncogenic Stemness Promotes Tumor-Sustaining Adaptability in the Intestinal Epithelium"

**Supplementary Materials**

ST1. Primary Antibodies used for immunohistochemistry.

| **Antibody** | **Dilution** | **Catalog#** | **Company** |
| --- | --- | --- | --- |
| Anti-CD44 Rat mAb, clone IM7 | 1:1000 | 103002 | BioLegend |
| Anti-GLS, clone 1D16 | 1:500 | ZRB1798 | EMD Millipore |
| Cdx2 (D11D10) Rabbit mAb | 1:1000 | 12306S | Cell Signaling Technology |
| Cleaved Notch1 (D3B8) Rabbit mAb | 1:500 | Val1744 | Cell Signaling Technology |
| Gfp (D5.1) XP® Rabbit mAb | 1:100 | 2956S | Cell Signaling Technology |
| Human/Mouse EphB2 Antibody | 1:100 | AF467-SP | R&D Systems |
| Keratin 20 (D9Z1Z) XP® Rabbit mAb | 1:1000 | 13063S | Cell Signaling Technology |
| Lysozyme, Polyclonal, Unconjugated, Ig fraction | 1:2000 | A009902-02 | Agilent Technologies |
| Smad4 (D3R4N) XP® Rabbit mAb | 1:250 | 46535T | Cell Signaling Technology |
| Tom20 (D8T4N) Rabbit mAb | 1:200 | 42406S | Cell Signaling Technology |

ST2. Secondary antibodies used for immunohistochemistry.

| **Antibody** | **Dilution** | **Catalog#** | **Company** |
| --- | --- | --- | --- |
| Goat anti-mouse IgG (H+L), Biotinylated | 1:300 | BA-9200 | Vector Laboratories |
| Goat anti-rabbit IgG (H+L), Biotinylated | 1:700 | BA-1000 | Vector Laboratories |
| Goat anti-rat IgG (H+L), Biotinylated | 1:300 | BA-9400 | Vector Laboratories |

ST3. Primary antibodies used for western blot.

| **Antibody** | **Dilution** | **Catalog#** | **Company** |
| --- | --- | --- | --- |
| beta Actin Antibody (C4) | 1:1000 | sc-47778 | Santa Cruz Biotechnology |
| Pink1 Antibody (38CT20.8.5) | 1:1000 | sc-517353 | Santa Cruz Biotechnology |
| PRDX3 Rabbit PolyAb | 1:2000 | 10664-1-AP | Proteintech |
| PRDX6 Rabbit PolyAb | 1:2000 | 13585-1-AP | Proteintech |

ST4. Secondary antibodies used for western blot.

| **Antibody** | **Dilution** | **Catalog#** | **Company** |
| --- | --- | --- | --- |
| m-IgGk BP-HRP | 1:2000 | sc-516102 | Santa Cruz Biotechnology |
| mouse anti-rabbit IgG-HRP | 1:2000 | sc-2357 | Santa Cruz Biotechnology |

ST5. Materials used for single-cell RNA-sequencing sample preparation.

| **Product** | **Catalog#** | **Company** |
| --- | --- | --- |
| 40-micron filter | 542040 | Greiner Bio-One |
| 6-well TC plate | 10861-554 | VWR |
| 70-micron filter | 431751 | Corning |
| Bovine SerumAlbumin | 5217 | Tocris Bioscience |
| DAPI | 5087410001 | Sigma-Aldrich |
| Dead Cell Removal Kit | 130-090-101 | Miltenyi Biotec |
| Dispase | 7913 | STEMCELL Technologies |
| DNase | 7900 | STEMCELL Technologies |
| EDTA | 324504 | EMD Millipore |
| FITC anti-mouse/human CD44 [IM7] | 103006 | BioLegend |
| PBS | BP399-4 | Fisher Scientific |

ST6. Materials used for organoid preparation.

| **Product** | **Catalog#** | **Company** |
| --- | --- | --- |
| Advanced DMEM/F-12 | 12634010 | Thermo Fisher Scientific |
| Animal-Free Recombinant Murine EGF | AF-315-09 | PeproTech |
| B27 Supplement w/o Vit A (50x) | 12587010 | Thermo Fisher Scientific |
| Corning® Matrigel® Basement Membrane Matrix, Phenol Red-Free, *LDEV-Free | 356237 | Discovery Labware |
| Glutmax 1, 100x | 35050061 | Thermo Fisher Scientific |
| HEPES | 15630080 | Thermo Fisher Scientific |
| N2 Supplement | 17502048 | Thermo Fisher Scientific |
| N-Acetyl-L-cysteine,cell culture tested, BioReagent | A9165 | Sigma-Aldrich |
| Penicillin Streptomycin sol | 97063-708 | VWR |
| Recombinant Human R-Spondin 1 Protein | 4645-RS-025 | R&D Systems |
| Recombinant Murine Noggin | 250-38 | PeproTech |

ST7. Other reagents and Materials.

| **Product Name** | **Catalog #** | **Company** |
| --- | --- | --- |
| ABC-HRP Vectastain kit | PK-4000 | Vector Laboratories |
| BCA Protein Assay Kit | 0023224 | Fisher Scientific |
| Hematoxylin | 26030-20 | Electron Microscopy Sciences |
| Hypoxyprobe Red549 Kit | 701 | Hypoxyprobe, Inc. |
| ImmPACT (TM) DAB HRP Substrate | SK-4105 | Vector Laboratories |
| Methyl green | ZH0804 | Vector Laboratories |
| NaCl | S271-1 | Fisher Scientific |
| NaF | A13019-30 | Alfa Aesar |
| Paraformaldehyde | 15714-S | Fisher Scientific |
| PMSF | P7626 | Sigma-Aldrich |
| Proteases inhibitor | P8340 | Sigma-Aldrich |
| Sodium Vanadate | 72060 | Sigma-Aldrich |
| Tamoxifen | T5648 | Sigma-Aldrich |
| Triton x-100 | 0694 | VWR |
| TRIzol | 15-596-018 | Thermo Fisher Scientific |
